# Creating a biomedical knowledge base by addressing GPT inaccurate responses and benchmarking context

**DOI:** 10.1101/2024.10.16.618663

**Authors:** S. Solomon Darnell, Rupert W. Overall, Andrea Guarracino, Vicenza Colonna, Flavia Villani, Erik Garrison, Arun Isaac, Priscilla Muli, Frederick Muriuki Muriithi, Alexander Kabui, Munyoki Kilyungi, Felix Lisso, Adrian Kibet, Brian Muhia, Harm Nijveen, Siamak Yousefi, David Ashbrook, Pengzhi Huang, G. Edward Suh, Muhammad Umar, Christopher Batten, Hao Chen, Śaunak Sen, Robert W. Williams, Pjotr Prins

## Abstract

We created GNQA, a generative pre-trained transformer (GPT) knowledge base driven by a performant retrieval augmented generation (RAG) with a focus on aging, dementia, Alzheimer’s and diabetes. We uploaded a corpus of three thousand peer reviewed publications on these topics into the RAG. To address concerns about inaccurate responses and GPT ‘hallucinations’, we implemented a context provenance tracking mechanism that enables researchers to validate responses against the original material and to get references to the original papers. To assess the effectiveness of contextual information we collected evaluations and feedback from both domain expert users and ‘citizen scientists’ on the relevance of GPT responses.

A key innovation of our study is automated evaluation by way of a RAG assessment system (RAGAS). RAGAS combines human expert assessment with AI-driven evaluation to measure the effectiveness of RAG systems. When evaluating the responses to their questions, human respondents give a “thumbs-up” 76% of the time. Meanwhile, RAGAS scores 90% on answer relevance on questions posed by experts. And when GPT-generates questions, RAGAS scores 74% on answer relevance. With RAGAS we created a benchmark that can be used to continuously assess the performance of our knowledge base.

Full GNQA functionality is embedded in the free GeneNetwork.org web service, an open-source system containing over 25 years of experimental data on model organisms and human. The code developed for this study is published under a free and open-source software license at https://git.genenetwork.org/gn-ai/tree/README.md.

## Introduction

One of the great challenges in biomedical research is the efficient discovery and summarization of rapidly expanding scientific findings. For example, as a conservative measure, MEDLINE, the National Library of Medicine’s bibliographic database on journal articles in life sciences, currently counts close to one million new publications per year [1]. Even on specific topics, such as diabetes or Alzheimer’s disease, no researcher can keep up with the rate of new contributions.

Here we present the GeneNetwork Question-Answer knowledge base (GNQA), a contextual generative pre-trained transformer (GPT) driven by retrieval augmented generation (RAG) for aging, dementia, Alzheimer’s and diabetes. GNQA builds on the GeneNetwork.org web service (see Fig. 1) — an open-source system containing over 25 years of experimental data on model organisms and human — and includes publications relevant to our biomedical research. As of this point GNQA incorporates one thousand publications that reference the GeneNetwork web service, along with one thousand research publications on human aging and one-thousand research publications on human diabetes.

**Figure 1.**
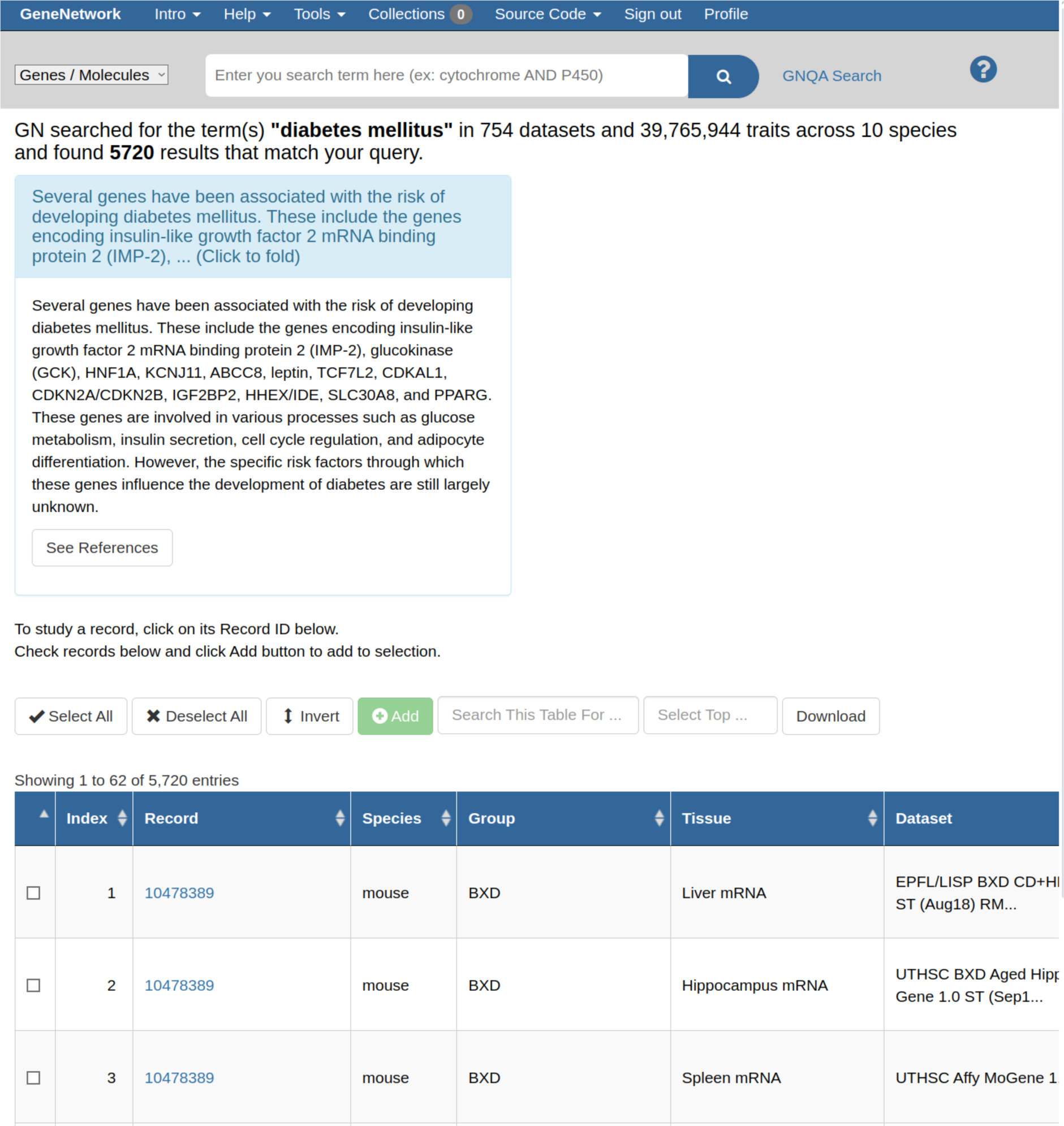
GNQA search is integrated in the genenetwork.org web-service (GN). GN is a web service that contains over 25 years of research data on model organisms. Our GNQA RAG augments and improves the service’s live search facility by mining information from thousands of manuscripts. The figure is a screen shot that showcases an answer in a small text box above the main GN search result table. The section in light-blue in which the answer is located has links to display the full answer, and a link that takes one to the standalone GNQA interface, where one can see the full answer and references to the exact sources, thereby providing full provenance.

**Figure 2.**
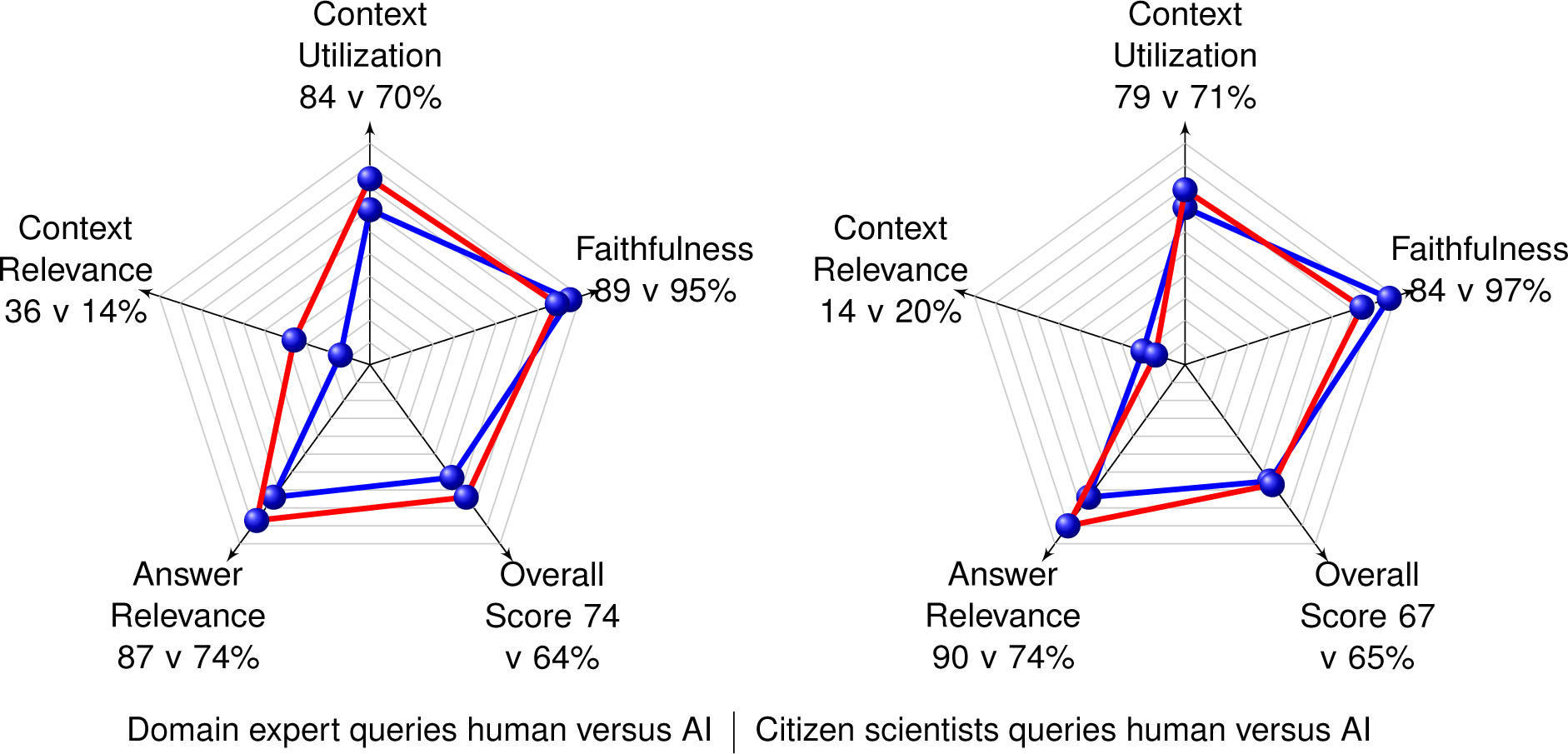
How well does AI generate questions for GNQA? As the usefulness of generative AI continues to improve, we look at how well it does at generating questions for our use case. In our experiment humans still generate better rated questions than AI. In the radar plot to the right, for citizen scientist questions, the red graph represents the RAGAS performance of humans while the blue represents RAGAS performance of AI generated questions. In the radar plot to the left, for domain experts, the red graph represents RAGAS performance of human questions while the blue represents RAGAS performance of a subset of the AI generated questions. It represents a subset as many of the domain expert AI generated questions could not be answered by GNQA. We rate the RAGAS scores that could be calculated. The questions generated by humans and AI have extremely similar scores, which shows the AI and the prompt for generating the questions are adequate and closely simulated non-experts. However, questions generated by domain experts have more obviously difference in score, a 10% difference, while only five out of 20 of the general questions generated by the AI produced measurable responses.

The last two years saw a marked increase in capability for generative AI, specifically with respect to natural language processing (NLP) powered by large language models (LLMs)[2, 3]. This class of GPTs have been found to answer questions, summarize information and are able to reason well enough to make users feel confident about responses[4]. Recent developments in NLP encouraged us to explore text search of scientific publications relevant to the topics of aging, dementia, Alzheimer’s and diabetes by parsing publications to build the GNQA and to evaluate our system with feedback from users. GNQA acts as a knowledge base that uses retrieval augmented generation (RAG) and leverages GPT. GPTs, now commonly available through ChatGPT, Claude, Gemini, etc., are a type of LLM, and the first GPT was introduced in 2018 by OpenAI[5] following Google’s introduction of the transformer architecture in 2017 [6].

A major issue with GPT, in the scientific context, is the injection of randomness, a generative aspect known as ‘hallucination’. A notable example is the fabrication of false references that have high apparent face validity. The problem is pervasive, and errors of commission by LLMs can include erroneous statements of facts, obvious reasoning errors, and other types of false positives[7, 8]. For our study, the goal is to provide provenance for AI-generated responses—to reduce errors of commission by retrieving contextual information and presenting original source data to users. This feature set enables users to compare the underlying text that led to the answer and, even better, the actual publications to validate contextual relevance.

RAG, introduced in 2020, is a retrieval-augmented technique capable of fetching information from a contained external database. Responses are limited to the data contained in that database thereby enhancing users’ confidence in the accuracy and relevance of LLM output[9]. In recent years, RAG technology has proven effective in the biomedical field. For example, Ge et al.[10] created Li Versa for liver disease queries, while Ranjit et al. applied RAG to radiology reports[11]. Yu et al. utilized RAG for diagnosing heart disease and sleep apnea[12, 13]. Recent advancements in RAG systems have showcased their transformative potential in various clinical and biomedical applications. Unlu et al.[14] utilized a RAG-enabled GPT-4 system to screen clinical trial candidates based on their electronic health records, comparing its efficacy to human performance over a two-year period without a direct assessment of the RAG system itself. Murugan et al.[15] implemented a RAG-based expert system to field pharmacogenomics (PGx) queries, benchmarking it against a specialized PGx catalog and GPT-3.5, highlighting its utility in personalized medicine. Bispecific Pairwise AI, detailed by another team, used a combination of GPT and XGBoost models with pairwise learning to select bispecific antibody target combinations where RAG was used to enhance the interpretability of ML model outputs, significantly improving the clarity of explanations for drug developers. Gue et al.[16] explored the efficiency of GPT-3.5 in extracting information from PubMed articles on diabetic retinopathy, comparing it to human extraction without employing a full RAG system, while Mashatian et al.[17] developed a RAG-based diabetes QA system, achieving 98% accuracy using the NIH National standards for Diabetes Self-Management education. Glicksberg et al.[18] applied zero-shot and few-shot RAG methods alongside Bio-Clinical BERT and XGBoost models to predict hospital admissions, demonstrating RAG’s comparable predictive power to trained ML models with the added benefit of providing explanations for its decisions. Additionally, Kresevic et al.[19] compared expert responses to RAG system outputs in interpreting hepatological clinical guidelines, further emphasizing RAG’s potential to enhance clinical decision-making. In contrast, Chen et al.[20] study on domain-specific LLM performance did not create or evaluate a RAG system but highlighted the limitations of generalized LLMs in clinical contexts. Overall, these studies underscore the promising capabilities of RAG systems in improving data interpretability, personalizing medical care, and supporting clinical decision-making. The accuracy of ChatGPT in diagnosing patients with primary and secondary glaucoma, using specific case examples, was similar to or better than senior ophthalmology residents. With further development, ChatGPT may have the potential to be used in clinical care settings, such as primary care offices, for triaging and in eye care clinical practices to provide objective and quick diagnoses of patients with glaucoma [21].

RAG effectively mines an authoritative knowledge base that lives ‘outside’ of the main LLM’s training data. RAG extends the already powerful capabilities of LLMs to specific domains or an organization’s internal knowledge base, all without the need to retrain the model. It is not only a cost-effective approach to improving LLM output such that it remains relevant, accurate, and useful in various contexts, but it also allows us to provide accurate references to the responses and those, in turn, help our users decide the usefulness of the response [9].

Once we had a responsive RAG with context and references, we wanted to benchmark how well that system worked. To assess performance we introduced the recently published retrieval augmented generation assessment system (RAGAS)[30] in combination with human feedback. Our study shows that GNQA unlocks relevant information from literature in the biological domains and is a knowledge base that complements the more standard database searches. In addition we created a framework that allows bench-marking AI generated questions and answers with context together with real user responses from domain experts and citizen scientists alike. RAGAS evaluates the effectiveness of RAG systems by requerying LLM responses and scoring for using multiple metrics, e.g. faithfulness, context utilization, context relevance, and answer relevance (see results section for more).

In the biomedical field, RAGAS was recently used by GastroBot: a Chinese gastrointestinal disease chatbot based on the retrieval-augmented generation. When evaluating GastroBot using the RAGAS framework the authors observed a context recall rate of 95%. The faithfulness to the source was estimated to be 93%. The relevance of answers exhibited a strong correlation with 92% [13].

We asked our users (domain experts and citizen scientists [22]) to submit a list of standard questions and then to pose their own. The responses were immediately rated by the users by “thumbs up/down” or “no answer”. We found that GNQA did slightly better with domain experts, probably because their questions were more relevant to the corpus and the answers therefore matched better. We generated datasets from the system’s responses to users’ questions and submitted them for RAGAS evaluation.

Next, we used GPT to generate questions. I.e. we used GPT to pose questions to GNQA [23] and assessed the responses with RAGAS. Interestingly GPT was closer to citizen scientists than to experts which suggests the generated questions were broader than the domain knowledge contained in the RAG database.

## Results

### A. GNQA provides a generative pre-trained transformer (GPT) knowledge base driven by a performant retrieval augmented generation (RAG) with a focus on aging, dementia, Alzheimer’s and diabetes

We started by feeding a corpus of three thousand publications to a RAG that utilizes OpenAI’s GPT for NLP. We included one thousand publications that refer to the GeneNetwork (GN) web service, a database that contains over 25 years of experimental data on model organisms, mostly on mouse and rat [24–29]. The aim was a GPT question-answer system for the users of the GN web-service that provides context based on GN’s search facilities. This corpus of publications forms the initial basis of our RAG system. The first set was a selection of one thousand papers that directly mention and used the GN online web-service. Because a few top research interests include aging, dementia, Alzheimer’s and diabetes we also added one thousand peer reviewed references each on the topics of aging and diabetes to the knowledge base, a total of three thousand reviewed publications.

GNQA uniquely provides provenance with the GPT responses by adding the titles of publications that were parsed and together formed the RAG answer/response (see Fig. 3). To make that possible, GNQA stores the context as the text of the research papers, or document, in one hundred word paragraphs. Each paragraph is therefore related to a document and stored in a database (Fig. 5). These paragraphs, which form the context. are part of the RAG inference information for a response. The user interface shows the answer (Fig. 3), followed by a list of references (Fig. 4) based on a *post hoc* RAG similarity search.

**Figure 3.**
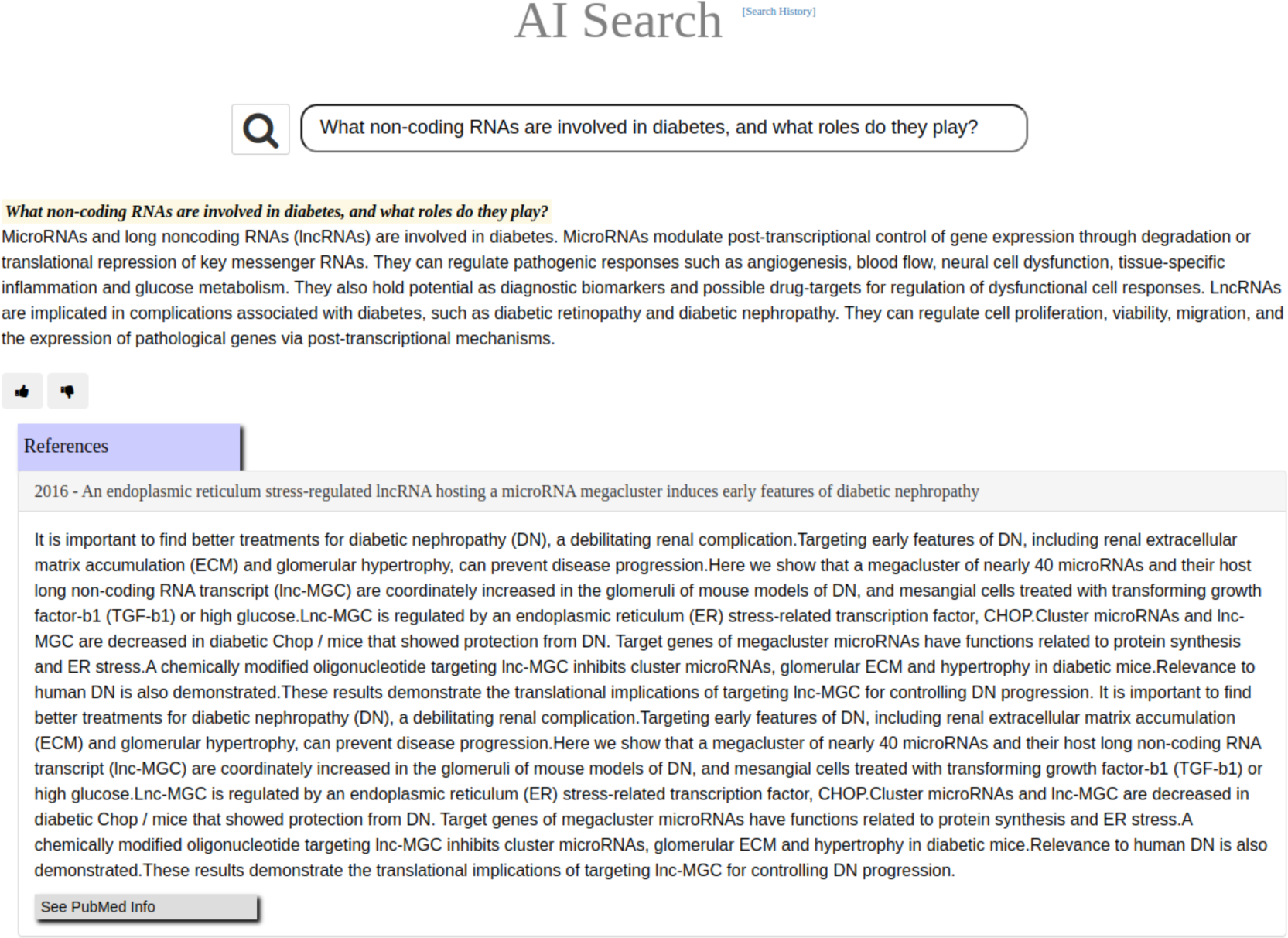
GNQA is a GPT-driven question-answer knowledge base, with provenance, for the GeneNetwork web-service. Here we show the response to the question: *What non-coding RNAs are involved in diabetes, and what roles do they play?* When questioning GNQA, the response is provided directly below the search bar. Following the response is a rating option, signified by a thumbs up and down, that allows users to provide immediate feedback on the system’s response. To provide full provenance the reference list is displayed to the research articles from which the answer was generated. In this case the textual context is displayed of a single paper on microRNA expression Fig. 6 (for a full list see Fig. 4). The listed references are notably not generated by GPT, but based on a RAG similarity search.

**Figure 4.**
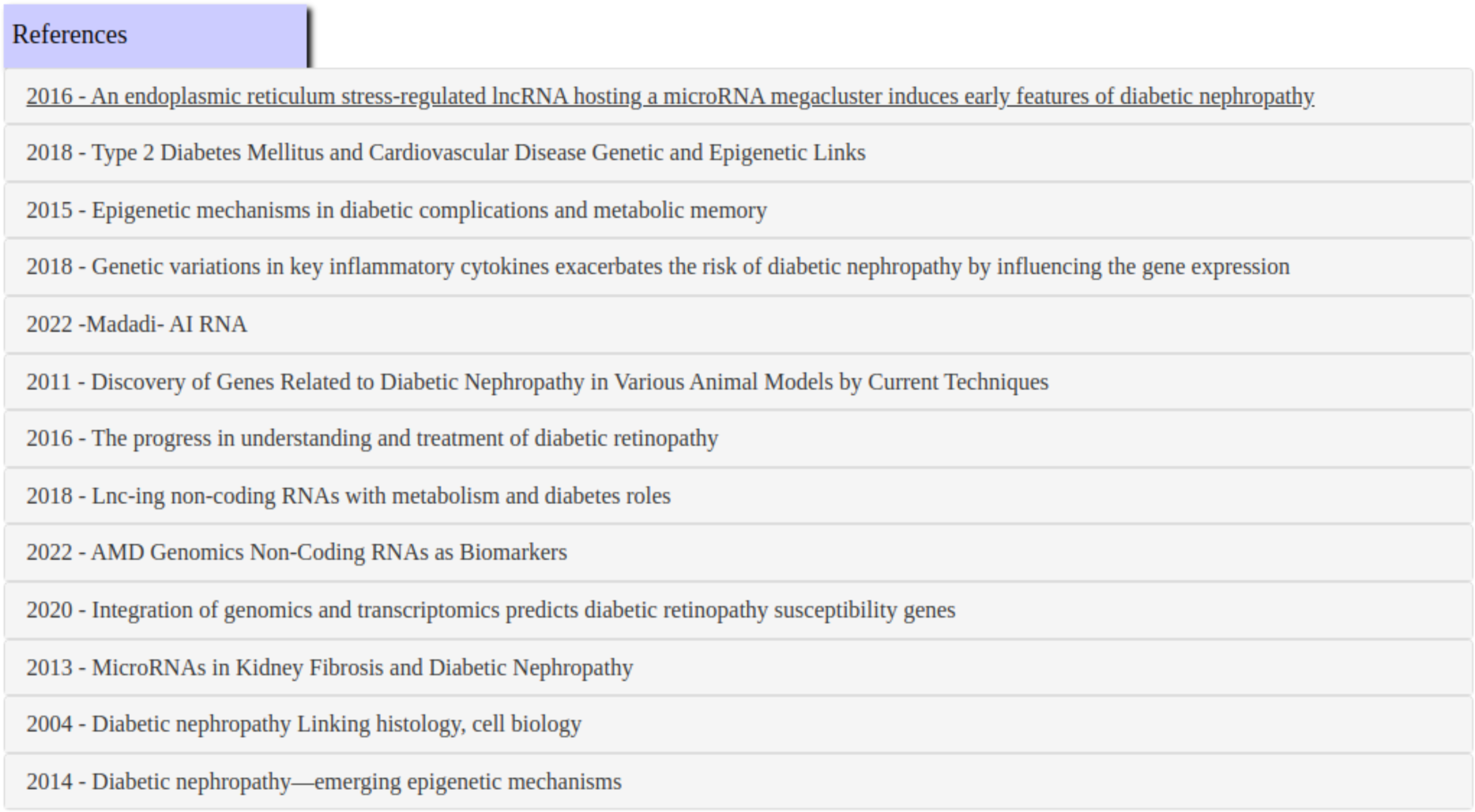
Provenance is provided by showing the full references list for the question: *What non-coding RNAs are involved in diabetes, and what roles do they play?* displays the titles of published research documents used for an answers. The references themselves are notably not generated by the system, but based on a RAG similarity search of the response. Clicking on a link will expand to the text that was used to find the answer (see Fig. 6 for an example).

**Figure 5.**
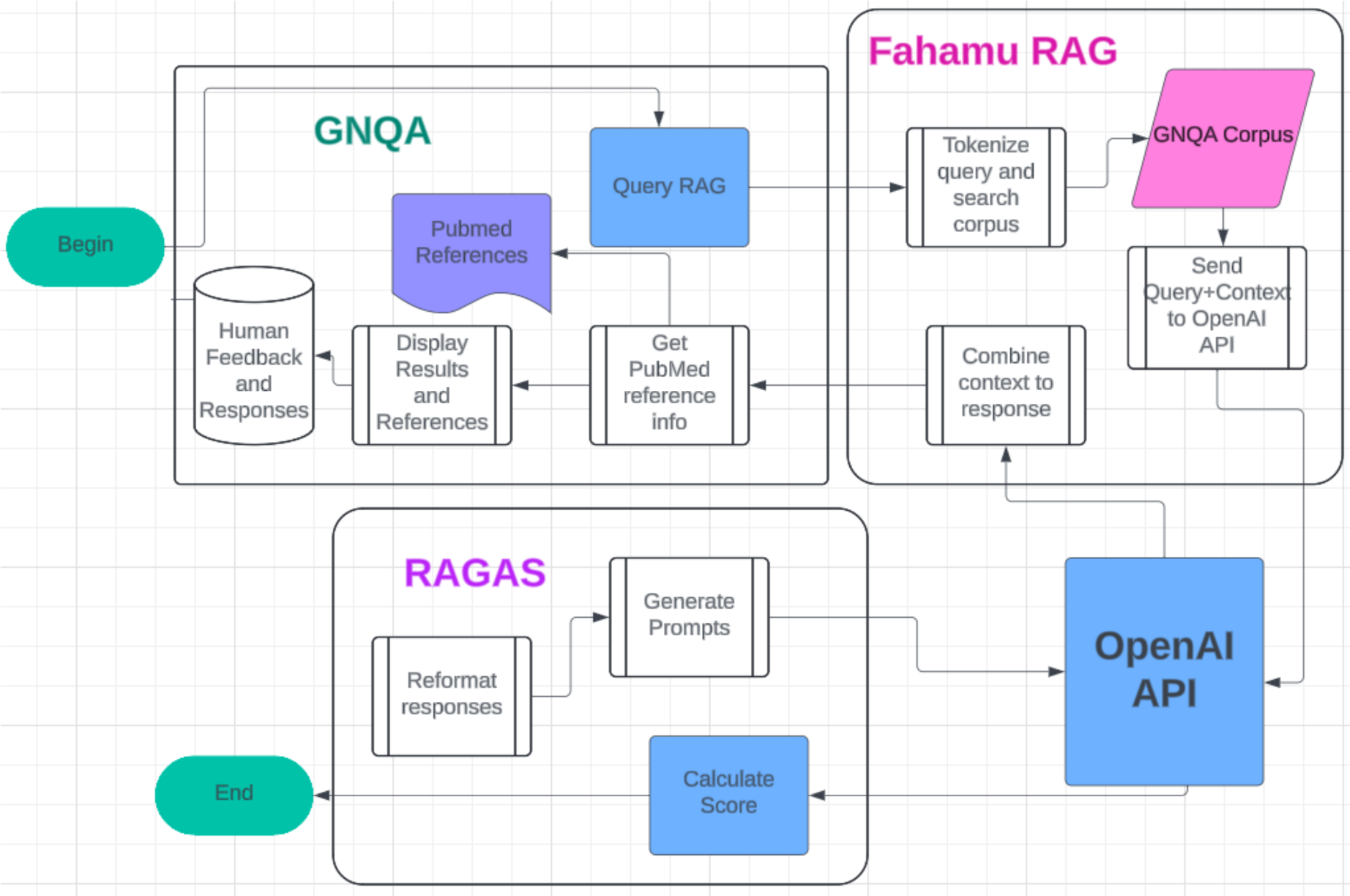
GNQA OpenAI RAG+RAGAS workflow. The main interaction point for the user is a web site that allows entering a question, query RAG, built by the GeneNetwork team. Three thousand documents on aging, dementia, Alzheimer’s and diabetes are stored as small sections of text as part of Fahamu AI’s RAG. Fahamu AI is a Kenyan startup founded by co-authors AK and BM. PubMed references are stored separately along with the GNQA interface. The question with the a subset of document sections are sent to OpenAI’s GPT which gives a direct response/answer that is fed back to the UI (see also Fig. 3). This answer is returned via the RAG, along with the document ids of the documents from which the sections are taken. The document ids are aligned with their titles, the titles are checked againts the PubMed data, and GNQA displays the answer with a list of references and available PubMed data and links. Next, to assess the correctness of GPT responses RAGAS checks the answers back against GPT and produces automated statistics that we compare with human interaction results.

A user question is answered by a set of paragraphs and every reference presents at least one paragraph in our database. Still, GNQA often finds the top matches for a question contain multiple paragraphs from the same research paper. GNQA collates and compiles these paragraphs into a single response text, which is added as an entity to the reference list and GNQA displays the compiled answer (Fig. 6).

**Figure 6.**
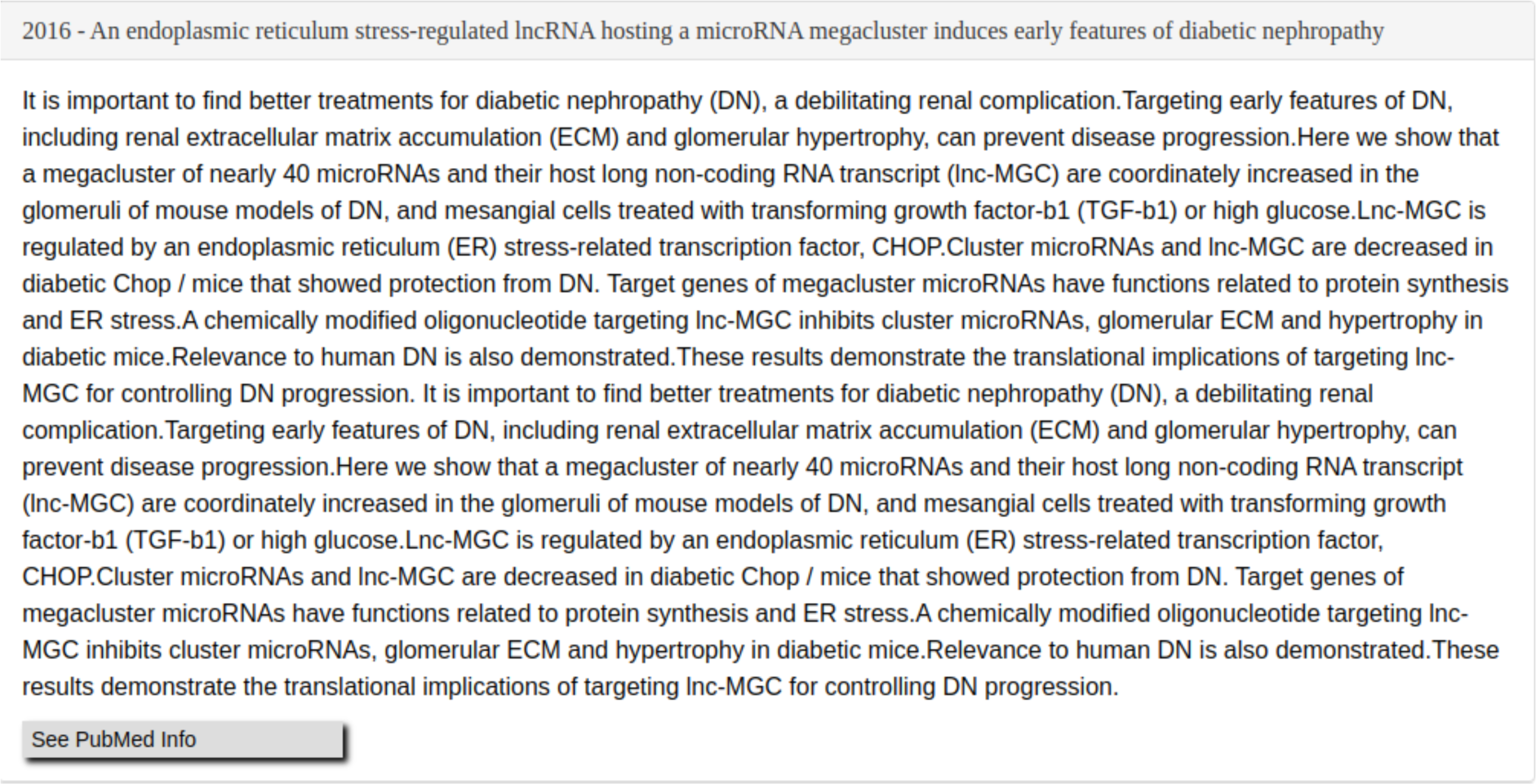
GNQA provides provenance to a response reference linked to the question *What non-coding RNAs are involved in diabetes, and what roles do they play?*. The response is followed by a list of publications (Fig. 4) and the text that formed the context for the answer. If multiple paragraphs from a single source are calculated as the top context entries, the textual paragraphs are combined as they are in this example.

Biomedical research papers are often available through NCBI’s PubMed [1]. PubMed has an open application programmer interface (API) that serves information about publications, including their abstracts, and web links to the hosted document[1]. When available, GNQA fetches references from PubMed’s API for the documents in its corpus, and stores it in the database to provide additional information on a referenced document and makes it findable online (Fig. 7).

**Figure 7.**
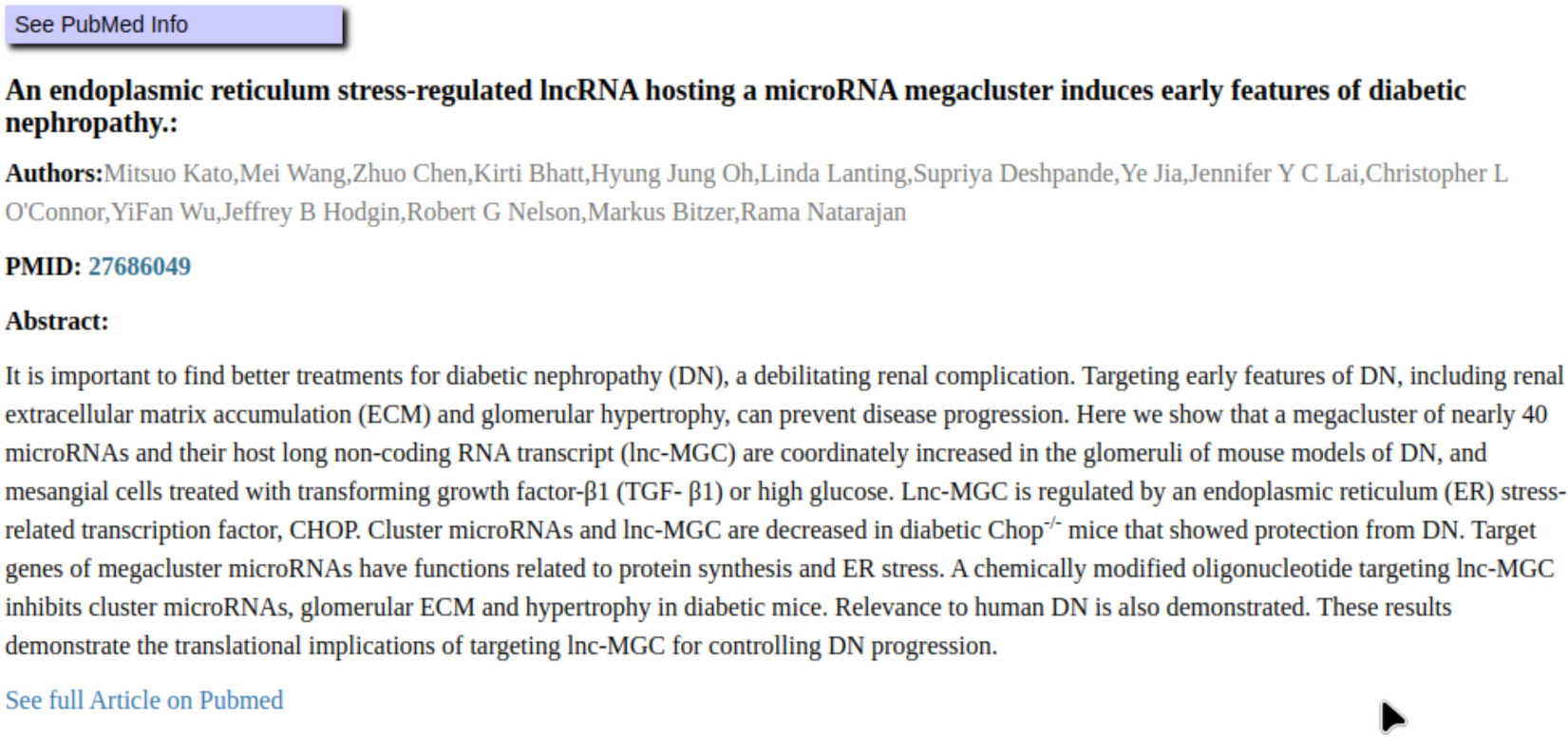
PubMed contains the full text of many of the references used by the GNQA. GNQA provides provenance through its references, and as NCBI’s PubMed is a widely used system for biomedical researchers, being able to get a link to a public reference makes responses more useful. For the question *What non-coding RNAs are involved in diabetes, and what roles do they play?* a reference is found through the PubMed API and the publication abstract along with a link to its full text is displayed.

We embedded GNQA into the main GN web service. On the main page of GN there is an NCBI-styled ‘global search bar’ with an example search term. A small resultbox has been added to the existing results interface that contains the full answer to a query generated for the keyword, and a link to the standalone GNQA results page with the answer and full reference list (Fig. 1).

### B. Performance rated by automated RAGAS evaluation

GPT not only can generate answers, it can also be used to evaluate results. **R**etrieval **A**ugmented **G**eneration **AS**sessment (RAGAS) is an automated tool that can evaluate RAGs[**Shahul:2023**]. We found that RAGAS provides useful measurements of relevance and faithfulness with respect to the questions, their context and returned answers. The following metrics: faithfulness, answer relevance, context relevance, and context utilization are all calculable with only the question, its answer, and the context from which the answer is derived by using secondary GPT queries[30]:

#### Faithfulness

Measures the factual consistency of the generated answer against the given context. It is calculated from answer and retrieved context. The generated answer is regarded as faithful if all the claims that are made in the answer can be inferred from the given context.

#### Context utilization

Context utilization is a metric that evaluates whether all of the answer relevant items present in the contexts are ranked higher or not. Ideally all the relevant chunks must appear at the top ranks. This metric is computed using the question, answer and the contexts, with values ranging between 0 and 1, where higher scores indicate better precision.

#### Context relevance

This metric gauges the relevancy of the retrieved context, calculated based on both the question and contexts. Ideally, the retrieved context should exclusively contain essential information to address the provided question.

#### Answer relevance

Assesses how pertinent the generated answer is to the given question. A lower score is assigned to answers that are incomplete or contain redundant information and higher scores indicate better relevancy. This metric is computed using the question, the context and the answer. An answer is deemed relevant when it directly and appropriately addresses the original question. Importantly, our assessment of answer relevance does not consider correctness instead penalizing cases where the answer lacks completeness or contains redundant details. To calculate this score, the LLM is prompted to generate an appropriate question for the generated answer multiple times, and the mean cosine similarity between these generated questions and the original question is measured. The underlying idea is that if the generated answer accurately addresses the initial question, the LLM should be able to generate questions from the answer that align with the original question.

We note that RAGAS offers additional metrics, but these require a known and correct answer for each question. For our free form queries these metrics are not usable[30]. For a full description of the RAGAS metrics used in this work see appendix D.3.

Table 1 shows that faithfulness is on average 87% across the selected topics and levels of expertise. Context utilization and answer relevance also have high values at average 81% and 89%, respectively. Context relevance/recall is the lowest value given from the assessment, below 24% on average, which gives credence to the strength of the models inference ability.

**Table 1.** Retrieval Augmented Generation Assessment (RAGAS) scores from GNQA for domain experts (left) and citizen scientists (right). We evaluated 52 unique questions asked by domain experts in the biological sciences (33 about general GeneNetwork.org, 6 about aging and 13 about diabetes). We also evaluated 61 unique questions that were asked by citizen scientists to GNQA (32 about general GeneNetwork.org, 13 about aging and 16 about diabetes). RAGAS was used to score on faithfulness, relevance, and utilization where ‘Faithfulness’ is a measure for a lack of hallucination, i.e. an answer that is purely faithful to the context has no hallucination. ‘Context utilization’ measures the overall degree to which the answer was pulled from the context list. We note that RAGAS scores questions from domain experts are higher than those from citizen scientists in all but one metric, answer relevance. ‘Context relevance’ is a measure of the information density of the context with respect to the question being asked. ‘Answer relevance’ measures the level to which an answer is addressing the question of interest.

| Research Topic | Faithfulness | Context Utilization | Context Relevance | Answer Relevance |
| --- | --- | --- | --- | --- |
| General | 88% vs 89% | 74% vs 68% | 14% vs 10% | 83% vs 81% |
| Aging | 87% vs 76% | 90% vs 91% | 62% vs 10% | 91% vs 95% |
| Diabetes | 97% vs 86% | 84% vs 78% | 24% vs 22% | 91% vs 93% |
| Topic Average | 90% vs 84% | 83% vs 79% | 33% vs 14% | 88% vs 90% |

### C. Performance rated by domain experts and citizen scientists

In addition to the automated RAGAS ratings above—an AI measuring the capability of an AI—we invited volunteers interested in the biological sciences to ask GNQA questions and provide ratings on the answers. Some of the volunteers decided to give additional feedback, a summary of which is provided (see result section E). A straightforward interactive rating system was employed for the evaluation by giving a thumbs up or a thumbs down to a response (see Fig. 3).

Calculation of GNQA response ratings use the following scheme: a thumbs up was given the value of 1, a thumbs down was given the value of 0, and a non-rating was given a value of 0.5. The average overall rating the volunteers give GNQA is a positive 77%. The volunteers question and rate GNQA an average of eight times. Instructions to the volunteers were to come up with and rate up to five questions of their own, and rating five questions they were given. Table 1 gives the RAGAS evaluation of the unique questions asked by humans. Volunteers submitted 29 unique questions about diabetes, 19 about aging, and 65 general systems genetics questions. Each question set was evaluated three times to obtain an average value. Table 1 presents the volunteers’ scores as a percentage, with domain expert scores to the left and citizen scientist scores to the right. Figure 2 gives the group average performance of GNQA when questioned by humans and evaluated by RAGAS: 74% overall score for domain experts versus 67% overall score for citizen scientists.

For experts the context relevance was higher by 19% on average. RAGAS includes a metric related to hallucination called faithfulness. One of the main purposes for building RAG systems is to reduce ‘hallucination’; therefore, the higher the faithfulness score, the better. GNQA is 6% more faithful when providing responses to domain experts rather than citizen scientists. Less resulting ‘hallucination’ for questions from experts could be due to their knowledge better matching what is stored in the RAG corpus, i.e., they ask questions closer to what is contained in the corpus. Multiple experts spoke to the need for the system corpus to include text books in addition to research papers. Answer relevance was close for experts and citizen scientists, with an average evaluation score of 88% for experts and 90% for citizen scientists. As noted with the evaluation results in table 1, one can see the system does better on the more focused topics of aging and diabetes, especially with respect to answer relevance, compared to general GN questions. An interesting finding is that the very simple thumbs up/thumbs down rating scheme ends up giving an overall score fairly close to the one from the automated analysis, i.e., 76% versus 70% (when averaging domain expert and citizen scientist overall scores from figure 2).

### D. Performance rated by RAGAS using GPT-generated queries

Above results point out that GPT is useful for evaluating and answering questions. Getting human involvement to evaluate an AI system is still necessary; however, more automation moving forward can increase the speed of development and evaluation.

Work by Argyle et al.[31] “demonstrates that these language models can be used prior to or in the absence of human data”. This led us to use the following prompt and queries to generate 120 questions for GNQA from the perspectives of a citizen scientist and expert, i.e., we asked GPT to respond either as a citizen scientist or an expert. For example:

There is a retrieval augmented generation system, called GNQA that holds a corpus of 3000 research documents. The documents span the topics of research related to genenetwork.org, research about the genetics and genomics of diabetes and aging. The systems topics will be referred to as GN, aging, and sugah. Two types of individuals question GNQA, citizen scientists and domain experts. A citizen scientist is someone with no more than undergraduate level understanding of biology and is someone who did not major or minor in biology. A domain expert has studied advanced biology and has a graduate degree in a type of biology or majored in biology for undergraduate school.

- Generate 20 questions, for GNQA about research on GN from the perspective of a citizen scientist.
- Generate 20 questions, for GNQA about research on GN from the perspective of a domain expert.
- Generate 20 questions, for GNQA about research on aging from the perspective of a citizen scientist.
- Generate 20 questions, for GNQA about research on aging from the perspective of a domain expert.
- Generate 20 questions, for GNQA about research on sugah from the perspective of a citizen scientist.
- Generate 20 questions, for GNQA about research on sugah from the perspective of a domain expert.

The system prompt the and questions were submitted to OpenAI’s GPT-4o. The machine generated queries we can compare with the human queries (see appendices for a list of all posed questions). For example, one result of this exercise is that the more focused domain topics of aging and diabetes make question generation easier for both humans and GPT. Another result is that GPT handled the concept of asking questions and the translation from ‘sugah’ to diabetes well. The majority of the automatically generated questions for the aging and diabetes topics received responses from GNQA; whereas, close to half of the generated general GeneNetwork systems genetics questions did not receive a response from GNQA. In a human citizen scientist vs. GPT comparison (see Fig. 2 and Table. 3) on citizen scientist questions and answer relevance, context utilization and overall score, humans score 16%, 8% and 2% higher than AI. In a human expert vs. GPT comparison (Fig. 2) human experts come up with quantitatively 10% better questions than GPT, and almost all of the human expert questions received answers from GNQA. Interestingly GPT generated questions on the topics of aging and diabetes were fully answered by the RAG. With GPT questions on GeneNetwork, however, results broke down with 75% of questions not answered (see appendix C.1). This is probably due to GPT splitting GeneNetwork into two words ‘gene network’. We note that the GPT questions are formulated from the wider OpenAI model and not from the RAG. That simulates the questions from humans coming out of ‘nowhere’ rather well. The scores in Table 2 and Fig. 2 for the automatically generated questions are based off of the questions that could be scored and we did not penalize the score for questions without answers. Qualitatively the automatically generated questions that were so out of bounds that GNQA could not properly respond to them suggests that we improve prompting (see appendix C.1). Additional possible explanations for GNQA’s inability to provide answers for some of the GPT-4o generated questions is that they are too creative for the system making it so the AI is actually asking questions beyond the realm of expert and/or the questions exceed the knowledge level gleaned from a three thousand publication corpus. I.e., too creative in that the biological knowledge the AI pulls from is so deep it has formulated questions that require a similar depth. The depth could be due to not only having knowledge of our corpus in biology, but of all the biological information that was been captured online to create the OpenAI LLM model.

**Table 2.**
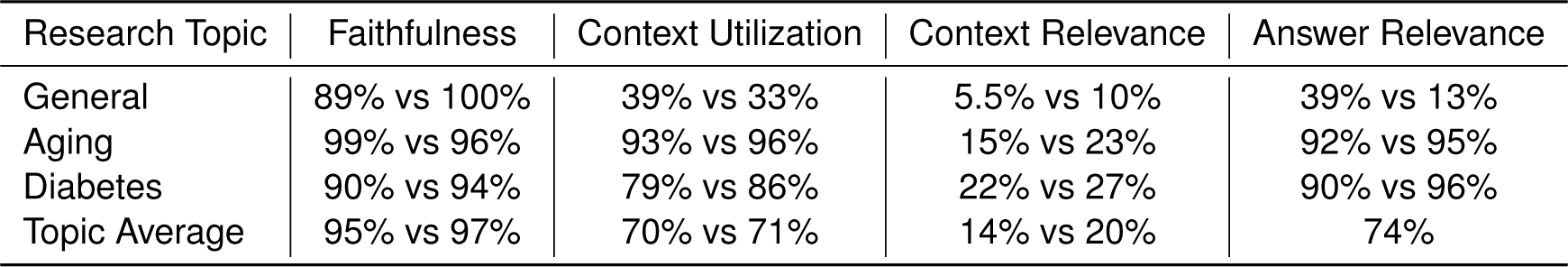
RAGAS results for GPT-4o generated questions to GNQA. We generated prompts through OpenAIs API playground resulting in 120 questions (see appendix – Questions). 20 questions were generated for each of the six areas out of the two levels of expertise, e.g. domain expert/citizen scientist, and three research topics: GeneNetwork.org research, aging and diabetes. The scores are shown as percentages, where the simulated domain expert scores are to the left in the column while the simulated citizen scientists scores are to the right. Unlike the trend with the human scores, the simulated domain experts do not perform better than the simulated citizen scientists based on RAGAS scores, but overall 2% worse. AI generated domain expert questions on diabetes and aging got responses, but most of GN related questions failed to get an answer from GNQA, possibly because the LLM split genenetwork into the words ‘gene network’.

**Table 3.** RAGAS scores for questions from human volunteers vs GPT4o generated. Human scores are to the left and GPT4o scores are to the right. The first set of values in the table are for questions from domain experts, human and GPT4o. The second set of values in the table are for questions from citizen scientists both human and GPT4o. For domain expert level questions the human volunteers scores are better in every metric except faithfulness. For citizen scientist level questions context utilization and answer relevance are higher than for the GPT4o generated questions.

| Domain Expert Scores Human vs GPT4o Questions |  |  |  |  |
| --- | --- | --- | --- | --- |
| Research Topic | Faithfulness | Context Utilization | Context Relevance | Answer Relevance |
| General | 88% vs 89% | 74% vs 39% | 14% vs 5.5% | 83% vs 39% |
| Aging | 87% vs 99% | 90% vs 93% | 62% vs 15% | 91% vs 92% |
| Diabetes | 97% vs 90% | 86% vs 79% | 24% vs 22% | 91% vs 90% |
| Topic Average | 90% vs 95% | 83% vs 70% | 33% vs 14% | 88% vs 74% |

**Table 3. RAGAS scores for questions from human volunteers vs GPT4o generated.**
| Citizen Scientist Scores Human vs GPT4o Questions |  |  |  |  |
| --- | --- | --- | --- | --- |
| Research Topic | Faithfulness | Context Utilization | Context Relevance | Answer Relevance |
| General | 89% vs 100% | 68% vs 33% | 10% vs 10% | 81% vs 13% |
| Aging | 76% vs 96% | 91% vs 96% | 10% vs 23% | 95% vs 95% |
| Diabetes | 86% vs 95% | 78% vs 86% | 22% vs 27% | 93% vs 96% |
| Topic Average | 84% vs 97% | 79% vs 71% | 14% vs 20% | 90% vs 74% |

### E. Overall user feedback

Informed expertise leads to better system testing. Half of the experts who tested GNQA provided additional feedback. These experts provide varying levels of expertise in the field, ranging from students (masters and doctorate) to full professors. Feedback from the doctoral students and professors was insightful as they are quite well read in the subject matter and asked questions directly geared toward their specific knowledge. By asking very specific questions they could evaluate reference relevance differently than RAGAS. Very specific questions refers to probing questions to let them understand the breadth and limitations of the corpus. Domain experts have knowledge of aging, dementia, Alzheimer’s and diabetes which enable them to ask questions that they may know the answers to or know which references are the best source for any possible answer to their question. RAGAS evaluates faithfulness, context relevance, context utilization and answer relevance without providing ‘correct’ answers alongside the question, answer, and context. Experts, although they are not familiar with RAGAS, include phrases and suggestions that speak to each of the four measurable RAGAS results, namely faithfulness, context utilization, context relevance and answer relevance.

#### An expanded evaluation

The domain experts rated responses using the following language: poor, OK, hard to rate (due to being partially correct or blatantly incorrect), incomplete, very good, passed reality check, not very helpful, closer to helpful, references no longer valid, good but high school level, B or C, and too focused on one part of the question. We found the experts had differing focuses, including improvement suggestions requiring differing interventions: additional references and tools, modified system prompting, similarity search modification, and incorporating agents. Upon reviewing the response from the following question ‘Which mouse genes have been associated with longevity?’ the expert gave the PMID (PubMed reference number) for a publication that was not in the reference list used as context for the answer to their question. According to our automated results GNQA performs best on the narrow topics of genetics/genomics with respect to diabetes and aging, and a major part of that performance is the lack of hallucination detected from its responses.

#### Improvement suggestion: deepen the corpus

The additional feedback from multiple experts suggests the need to add fundamental works on the subject matter to GNQA, as in adding more than just research papers. Improving upon the state of the art is often incremental; hence, requiring multiple steps or stages to be taken. A software system is being built with curated data, advanced artificial intelligence, FAIR standards to work in conjunction with an existing systems genetics tool. As a first step our expert system is an advanced question-answer system with thousands of research documents germane to GN, the genomics of aging and the genomics of diabetes. It was initially believed many research publications was a great start; however, after receiving feedback from experts, we find the experts expect GNQA to be more well versed in general genomics and genetics, not confined to the topics most closely related to the topics in our corpus. GNQA needs more information, perhaps even wikipedia and/or books on biology, advanced biology, genetics, and genomics added to its corpus to better meet the knowledge level of an expert. Beyond adding books to GNQA’s corpus, five experts brought up the fact that the knowledge base is missing several important tool references and publications, while containing some outdated tools references. Positive feedback was that, because the corps contained information an human and model organisms, it was generating interesting translational responses. Discovering information that translates to mouse to human and *vice versa* is one of the grand challenges in biomedical model organism research.

Discovering datasets hosted in GN on a topic of interest is one of the main goals of building GNQA. E.g. a user coming into GN and looking for diabetes has a chance of finding relevant experimental model organism data. GNQA adds value because it gives information on relevant contextual information, including our corpus of manuscripts uploaded into the RAG. Adding additional information to the corpus, to provide more context, is the easiest and quickest improvement suggestion to implement. We are currently in the process of doubling the corpus by uploading more publications. Another possibility would be to glean more from the underlying GPT system (currently OpenAI) and include that as additional ‘hints’ with a notification of the possibility of hallucination. Three experts brought up that it matters in what way a question is worded — and them being able to influence and improve the response. This suggestion shows that a savvy user can quickly adapt question formation to optimize output, similar to how we enhance google queries, but this is probably hard without *a priori* domain knowledge and expertise. It would be even better if, in the future, GNQA can adapt to a user’s style of questioning.

#### Improvement suggestion: better LLM prompts

This second main improvement suggestion requires modified prompting, and may be solved by introducing a prompt into the workflow to rephrase the question, then provide multiple answers whose efficacy can be rated by the user. We would have to incorporate the feedback into GNQA at runtime to allow adaptation. Without implementing any form of model re-training, a workflow modification can flip a flag for the user that turns on alternate questions, while saving in the user’s profile a description of the types of question re-framing that best supports response correctness. As researchers we need to figure out how to describe one’s re-phrasing preferences to accomplish this goal. Three experts spoke explicitly to references not being proper context for an answer.

#### Improvement suggestion: enable agents

Improvement suggestion group three leads to the idea of using agents. Agents will query different application programming interfaces (APIs) to get up-to-date data to provide a truly best possible inferred response, an answer with context. For example, research is published on a daily basis, and if there was an agent that could query an API for new publications and use those as possible context to answer questions, it would improve answers and ensure an up-to-date reference list. Another agent could question ChatGPT, retrieve its answer, check how much of that context can be found in GNQA’s complete context corpus then incorporate all data into a response. A major strength of GNQA we have developed is the high fidelity of its references, and maintaining provenance with every response. As ChatGPT now cites other websites to show a shallow level of provenance, another agent could query these to retrieve concrete references to very specific documents, further improving responses.

## Discussion

One of the great challenges in biomedical research is unlocking published information for researchers in an accessible way. Even on specific topics, such as diabetes or Alzheimer’s, it is impossible for any researcher to keep up with the rate of produced scientific manuscripts. NLP has recently turned a corner after the introduction of transformers[6]. Transformers have helped scale up LLMs, thereby improving NLP, so that they now closely appear to mirror human communication. Recent studies have demonstrated the ability of LLMs to process and generate information, exemplified by OpenAI’s ChatGPT[5], Google’s Bard [32], Facebook’s Llama [33, 34], and Databrick’s Dolly[35]. These models have been applied in biology where they have shown promise in tasks such as literature review and information retrieval[36–38]. We believe application of LLM on research literature in conjunction with databases and their metadata may be one of the most useful applications of NLP for the biomedical sciences. To explore LLMs we started by mining a small set of literature in a small number of topics.

One of the main limitations of current LLMs is that responses lack the context and specificity required for scientific pursuit[39]. A major issue in the scientific context is the generative aspect of GPT, also known as ‘hallucination’. GPT suggests fictional facts and makes reasoning errors, lacks up-to-date information, and has confidence in false positives, which is particularly alarming for use in public health[7, 8]. GPT’s usefulness therefore presents a problem: it lulls people into a dangerous complacency. When AI systems are very good, people tend to trust the output without fully scrutinising it. When the AI is good but not great, people are more attentive and add their own judgment, which improves performance. It is a reminder that in this new age of AI humans are still needed — and humans must still be sharp and verify responses.

We introduced a RAG on top of OpenAI’s API and added three thousand peer-reviewed documents on aging, dementia, Alzheimer’s and diabetes. For our study the goal for the GNQA was to provide provenance for its responses, i.e. after getting the first response, the RAG searches again and fetches contextual information and presents that to the user (see Fig. 5). These references allow users to visit and rate the underlying text that led to the GPT response and, even better, references to actual publications to validate contextual relevance. To assess performance of our knowledge base, we collected human feedback from domain experts and citizen scientists and introduced automated RAGAS assessment that uses GPT to validate responses. We asked our users — consisting of biologist domain experts and ‘non-biologist’ citizen scientists — to submit a list of standard questions and then to pose their own questions. The GPT responses were immediately rated in the web interface by the users by “thumbs up/down” and “no answer”. From the human feedback we found that GNQA performed slightly better with domain experts. This suggests expert questions are more relevant to the corpus and therefore generates a better matching GPT response. Next we submitted the same responses to RAGAS for evaluation. Of the values RAGAS measures, ‘answer relevance’ is the closest to our thumbs up/down response relevance, which considers, not only the answer, but also the context or references returned with it. RAGAS machine evaluation of ‘answer relevance’ has an average score of 89% versus 76% for the human thumbs up/down user rating. This points out that in our setup RAGAS is more positive about GPT response than humans and we conclude that RAGAS can not completely replace humans for assessment. Even so, RAGAS is a useful automated assessment system which we can use to check and fine tune our system down the line.

Next, we asked GPT to generate questions from our knowledge base and we used RAGAS to assess the response. Interestingly, the type of GPT generated questions scored closer to citizen scientists than to experts. It suggests the GPT generated questions were wider, i.e., less focused on the domain knowledge in our knowledge base. This result encourages us to keep pursuing automated generation of questions to test different LLM inference back-ends. In the future, if we get the resources to build our own models, we can use RAGAS as a goal function to reinforce and improve the GPT generated ‘domain expert’ questions.

It is important to note that the use of LLMs in the biomedical sciences raises ethical and societal concerns. GPT-generated text in scientific publications raises questions about the role of human authors and the potential for misuse or manipulation of scientific information[40–45]. The use of AI models to generate answers to biological questions raises concerns about the potential for bias in the training data and the impact on the reproducibility of scientific research [46–48]. With GNQA we allow scientists to find the information that led to the GPT response and that gives them a chance to assess the relevance and accuracy of the response. We introduced continuous human feedback from users as an important feature of our knowledge base. In addition, independent, accurate, reliable benchmarks for AI models matter. Benchmarks “define and drive progress”, telling model makers where they stand and incentivising them to improve. Benchmarks chart the field’s overall progress and show how AI systems compare with humans at specific tasks. Benchmarks are expensive to develop, because they often require human experts to create a detailed set of questions and answers. Automated GPT generated questions may help improve the system, but it also may introduce bias because the questions are derived from the GPT mechanism itself. In this study we had humans generate questions, we used GPT to generate responses and we used human feedback to assess the response. Separately we used GPT to generate questions, we use GPT to generate responses and we use GPT to assess the results. The GN web-service is a FAIR[49, 50] data resource and all of its source code is free and open source software[24–29]. Unfortunately, at this point, most LLMs and even their APIs are non-FAIR and closed source. Another limitation of current LLMs is their reliance on an unknown corpus of input data, pre-trained models, and a lack of transparency in their training data. While open-source models such as Dolly by Databricks and Llama by Meta have made it possible for researchers to extend and fine-tune the models for specific use-cases, the weights of the models are still kept confidential[34, 35]. For this study and building the user interface and RAGAS benchmarking system we pragmatically opted for OpenAI’s proprietary GPT and FahamuAI’s RAG solution as a service. This provides a base line for future work with free and open source software solutions when they appear. Our GNQA front-facing code is open-source, and the code used for evaluation — including RAGAS and bespoke code for managing data from GNQA — is also open-source [36–38]. In the next phase we will experiment with open source LLM models and use our benchmarks to compare closed source solutions with upcoming free and open source software solutions. We aim to introduce an open source LLM, such as Llama, Mamba or Olmo — a recent open source LLM that includes open training data [51] — and introduce an open source RAG engine, such as R2R [52]. Future work will also include working on the performance aspects of RAG solutions. Non-linear scalability is a topic that comes up in RAG discussions [53] and for a growing corpus of manuscripts retrieving fast responses is key for satisfactory user interface development.

In conclusion, GNQA now acts as a knowledge base with contextual information and is part of the GN web service[24–29]. GNQA complements the more standard database searches of GN. In addition we created a usable framework that allows bench-marking AI generated questions and answers together with real user responses from domain experts and citizen scientists alike. Our contextual knowledge base, aimed at aging, dementia, Alzheimer’s and diabetes, is uniquely designed and fine-tuned with quantitative automated and qualitative user feedback, making its evaluation a novelty among current advanced biology retrieval augmented generation (RAG) projects. We note that recent other publications on LLMs in biology did not incorporate user feedback[36–38, 54] except Zhou et al.[13] who performed a system usability evaluation. We also note that OPENAI has recently introduced a cursory provenance functionality by providing site references for responses[55]. The level of specificity of this service, however, is too low for scientific pursuit, as OPENAI will reference the publisher — including PubMed, Nature, or another scientific publisher — but not the actual text, nor a specific single document reference. This high-level GPT functionality does currently not compare with the fidelity of our RAG, though it may get there in the future.

## Methods

Implementation of GNQA involves data curation and ancillary tools in addition to taking advantage of a state-of-the-art LLM. GNQA interacts with OpenAI’s GPT-4 and FahamuAI’s “digest” through an application programmer interface (API) and organize the curated data into a graph structure that supports fast data retrieval (see Fig. 5). FahamuAI is an AI startup with a product called “digest” that streamlines the deployment of state-of-the-art question and answer systems. The GNQA web-interface is built as a python flask application to fit the style of GeneNetwork.org and the source code is available online. As a proof-of-concept implementation, functionality is limited to the base requirement of a question and answer system, and will undergo a user interface evaluation and redesign in later iterations, alongside a fine-tuning of the systems curated data, question prompts, and responses. Drilling down further into implementation methods we explain document acquisition, document ingestion, retrieval augmented generation (RAG), and our pursuit of GNQA being findable, accessible, interoperable and reusable (FAIR) [49, 50]. GNQA’s workflow and data flow through its main connected services can be found in Fig. 5.

### A. Data Acquisition

Our prototype is currently comprised of data from three domains: publications mentioning GeneNetwork.org, genetics/genomics of aging publications, and genetics/genomics of diabetes, and GeneNetwork.org metadata. A thousand research documents, in portable document format (PDF), each on GN, the genetics/genomics of aging, and the genetics/genomics of diabetes were collected and supplied to GNQA as curated context for questions.

### B. Document Ingestion

The text is extracted from portable document files, and that text is transformed using the GROBID [56] machine learning library for extraction, parsing and re-structuring of raw or unstructured documents. GROBID transforms the document text into structured descriptive XML/TEI (eXtensible markup language/text encoding initiative) documents. GROBID is a library that focuses on the transformation of technical and scientific articles to support their machine readability. GROBID enables natural language processing and other AI tasks to be done on previously unstructured data. The XML/TEI chunks are then made to fit within word length windows with approximately 100 words, the restructure respects the boundaries of sentences (to avoid bad context as much as possible). Resultant word chunks are stored in special PostgreSQL record types.

### C. Retrieval Augmented Generation (RAG)

We use hierarchical navigable small world graphs (HNSW) [57] as it has state-of-the-art performance on similarity search. HNSW graphs allow nearest neighbor search in logarithmic time without the need for additional data structures. GPT3.5/4/4o are used as a zero-shot binary classifier, as it is asked whether or not a returned paragraph answers the posed question. GPT3.5/4/4o return a logarithmic probability along with a ‘yes/no’ which provides a numeric way to rank the returned word chunks. The best chunks plus the question are sent to GPT3.5/4/4o to generate an answer, while the best word chunks are listed as references and are returned with their document titles. In the case of multiple word chunks coming from the same source, the word chunks are concatenated in the returned references. This ensures unique reference entries being presented with GNQAs answer to the question.

### D. RAGAS

RAGAS software requires a dataset to be in a specific format where questions, answers and context lists are themselves in lists. Datasets are designed to be evaluated using OpenAI endpoints and evaluating datasets with more than seven entries can cause issues with OpenAI request timing limits; hence, the datasets have between five and six entries. The output of the ragas evaluation is a JSON object with the metrics as keys, in this evaluation the metrics are faithfulness, context utilization, context relevancy, and answer relevancy. Each dataset was evaluated three times and the final values were averaged over the runs. Each set of questions is evaluated in groups of five; therefore, the averages for multiple datasets with the same topic is necessary. The other code and data files are used for preparing data output from the study and RAG endpoint.

### E. Data FAIRness

A FAIR system must follow its principles, summarized by: findable, accessible, interoperable and reusable. GNQA is part of GN which is findable and its main functionality is accessible to anyone. According to Mulligan *et al.* [28], GN is a tool for studying covariation and causal connections among traits and DNA variants in model organisms. GN is free open source software (FOSS) that uses community standard tools, protocols, and databases. All GN’s data is published as FAIR data[24–29].

In this era of LLMs, code and models, i.e., data, are often covered by the same licenses. Meta owns and has released a state-of-the-art LLM Llama2, recently Llama3, and its code and model are published under the same Llama community software license that basically asserts Meta’s intellectual property [58]. Huggingface.co [59] is a machine learning site that is a well-known repository for datasets used in the creation of state-of-the-art models is released under a more liberal Apache2 license [60] that allows usage of code and models as long as proper attribution is maintained in reproduction, sharing and usage. Machine learning models on Huggingface have “model cards”. A Huggingface model card is an identity document edited by the ML model contributor who then assigns a specific license for the model or dataset which can differ from that of Huggingface. For our work on GN and GNQA we use the web-service Affero GPL[61] for our code and corpus.

With LLMs there is the ongoing discussion about intellectual property (IP) of the learning data. For example, unwittingly, we also may be using publications and data picked up from the internet and considered ‘private’ or IP restricted. Part of the effort will be going into making sure there is proper attribution where required and filtering of data and the focus on provenance will help bring out and resolve any issues. With provenance we track publications and their sources and we will make sure that people can only access them with the relevant (journal) permissions.

### F. Volunteers

Volunteers are split into two types of individuals who evaluated GNQA: citizen scientists and domain experts. A citizen scientist is someone who did not major nor minor in biology. A domain expert has studied advanced biology and has a graduate degree in a type of biology or majored in biology for undergraduate school. Volunteers were solicited from the GN community; including professors, postdocs, doctoral students, software engineers and vendors. Volunteer results are based on feedback from eleven domain experts and ten citizen scientists.

### G. Data and source code availability

All questions, both posed by users and generated by GPT, are listed in the appendix. RAGAS evaluation was carried out using open source software. Examples of datasets that are ready to be evaluated by RAGAS are found in our source code repository.

## Funding

The authors gratefully acknowledge support from National Institutes of Health/NIDA U01DA047638 (EG and HC), National Institutes of Health/NIGMS R01GM123489 (PP and EG), and NSF PPoSS Award #2118709 (SD, PP, AG, CB, PH, ES and EG), and the Center for Integrative and Translational Genomics (EG).

## Acknowledgements

The authors thank the individuals who tested the system and shared their insightful discussion and feedback, and OpenAI/GPT for providing an easy to use API to their systems.

## Conflict of Interest

A. Kibet and B. Muhia are founders of FahamuAI, a company that provided a RAG service and API used in this study. The remaining authors declare no competing interests.

## Appendix - Questions

## A. Domain Expert Questions

### A.1. General

1. What are the potential benefits and risk associated with gene editing technologies like CRISPRR-Cas9?
2. How does epigenetics inluence gene expression without changing the underlying DNA sequence?
3. Describe the role of mitochondrial DNA in heredity and how it differs from nuclear DNA.
4. What are the ethical considerations surrounding prenatal genetic testing and the selective termination of pregnancies based on genetic factors?
5. Create a how-to guide for genetic sequencing.
6. Which genes give a predisposition to developing T1D?
7. What is ensembl
8. Which database can I use for genetic, genomics, phenotype, and disease-related data generated from rat research?
9. What is RGD?
10. What resources can I use to do pathway analyses?
11. Once a sperm combines with an egg, what determines how traits are passed onto the resulting lifeform?
12. Why is genetic tracing matrilineal rather than patrilineal?
13. Explain the process of DNA replication and how it ensures accurate copying of genetic information during cell division.
14. What are the potential benefits and risks associated with gene editing technologies like CRISPR-Cas9?
15. How does one tell the difference between X and Y DNA, with repsect to DNA tracing and determining QTLs?
16. For text and biological resources, do you mean add some books (on biology stuff) or/and web resources (as ensembl) on your system?
17. what is ensembl?
18. What is the difference between QTL mapping and GWAS?
19. How do I determine which gene in my QTL is causal for the trait?
20. Why do males have two Y chromosomes and females only one?
21. How does one tell the difference between X and Y DNA, with respect to DNA tracing and determining QTLs
22. Once a sperm combines with an egg, what determines how traits are passed onto the resulting lifeform?
23. How can I add a new species to the GeneNetwork database?
24. which genes are typically associated with diabetes in QTL analyses?
25. In which diseases is the gene TCF7L2 involved?
26. Once a sperm combines with an egg, what determines how traits are passed onto the resulting lifeform?
27. Can you explain what a ribosomal binding site at a high level and make it accessable to a non-expert?
28. Once a sperm combines with an egg, what determines how traits are passed onto the resulting lifeform?
29. Can you explain the difference between sequencing with short reads vs long reads? Please make you answer accessible to a non-expert.
30. Can you explain why using a pangenome-based reference might be more useful than simply using a single linear reference? Please make you answer accessible to a non-expert.
31. Is all genetic regulation done through DNA (e.g., prompters, repressors, activators) or are there other forms of genetic regulation? Please make you answer accessible to a non-expert.
32. What are the different relationship between traits?
33. Can landscape of QTL and GWAS hits be used to find relationships between traits?

### A.2. Aging

1. What is the significance of the length of telomeres?
2. Which mouse genes have been associated with longevity?
3. what genetic factor are associated with aging
4. which genes are typically associated with early aging?
5. How do I generate a linkage or association mapping study in mice to understand aging?
6. is there a specific genetic variation that can cause someone to live longer? please make your answer accessible to a non-expert

### A.3. Diabetes

1. How is gene expression in the liver affected by diabetes?
2. Is any of the genes SH2B3, IFIH1 or ERBB3 related to diabetes?
3. nutrition is a factor for diabetes. how can genomics be use to better understand nutritional factors of diabets
4. nutrition is a factor for diabetes. construct an abstract about how can genomics be use to better understand nutritional factors of diabets
5. Is the gene TCF7L2 involved in diabetes?
6. Is any of the genes SH2B3, IFIH1 or ERBB3 related to diabetes?
7. How can I use genenetwork to find genes related with diabetes in humans?
8. How can I use the GeneNetwork tool to find genes related with diabetes in humans?
9. what are confounding factors in diabetes?
10. How is the immune system related to diabetes?
11. What are the genomic variants associated with immune system components and diabetes?
12. What is the role of the immune system in the metabolomics of diabetes and associated conditions?
13. Can the landscape of QTL and GWAS hits be used to dissect the role of immune system in diabetes and complications?

## B. Citizen Scientist Queries

### B.1. General

1. What is the most cited environmental factor for the onset of asthma?
2. How would one extract the DNA, from say, flora or fauna?
3. genetics
4. what is bioinformatics
5. Explain the process of finding a genetic marker followed by a quantitative trait loci.
6. What about recombination in human centromeres?
7. How does recombination work in human centromeres?
8. What about recombination in the human genome?
9. Create a how to guide for genetic sequencing
10. What is the significance of the length of telomeres?
11. Once a sperm combines with an egg, what determines how traits are passed on to the resulting lifeform? ”,
12. Why is genetic tracing matrilineal rather than patrilineal? ”,
13. How does one tell the difference between X and Y DNA, with respect to DNA tracing and determining QTLs?”,
14. what type of dataset is useful for qtl mapping analysis in genenetwork2? ”,
15. what are the bioinformatics tools for QTLs analysis?
16. what are the statistical approaches for qtls analysis?
17. Create a how-to guide for GWAS analysis?
18. Create a how-to guide for genetic sequencing
19. Create a how-to guide for genetic sequencing.
20. What is the significance of the length of telomeres?
21. Create a how-to guide for genetic sequencing
22. Create a guide for genetic sequencing
23. Define dyslipidemia.
24. What is cytochrome?
25. How does one tell the difference between X and Y DNA, with respect to DNA tracing and determining QTLs?
26. how does environment influence fertilisation
27. how does diet impact someone’s height
28. which animal has the same number of chromosomes as human
29. what’s ensures brains work
30. how do our brains maintain emotions
31. what hormones do our brains release during stressful experiences?
32. what is the use of corticosterone?

### B.2. Aging

1. List as many studies as you can that include rapamycin.
2. Why is it so diffuclut to map gene loci that control aging in humans?
3. What is apoptosis?
4. which genes are involved in the aging process
5. what causes the aging process
6. which genes are involved in aging
7. what genes are involved in the aging process
8. Describe the genotypes related to Alzheimers and dementia which have commonalities with those for aging.
9. Describe the genotypes related to Alzheimer’s and dementia which have commonalities with those for aging.
10. What genetic factors influence aging in humans?
11. what genes are associated with aging?
12. Which genes are associated with aging in human
13. What is GeneNetwork and how does it relate to aging research?

### B.3. Diabetes

1. What are the genetic bases for the varying efficacy of diabetes treatments among individuals?
2. Explain Protective Genetic Factors Against Diabetes in Elderly Populations
3. Explain Effect of Lifestyle Modifications on Aging-Associated Diabetes Risk
4. Explain The Role of Longevity Genes in Protecting Against Diabetes
5. What are the types of diabetes
6. How many types of diabetes exist?
7. Is there a direct association between aging and susceptibility to having diabetes?
8. How does genetics influence the emergency of diabetes?
9. what genes are associated with diabetes?
10. What causes diabetes?
11. Does cycling reduce risk of diabetes?
12. How can GeneNetwork assist in identifying genetic factors involved in diabetes?
13. What specific tools within GeneNetwork are most useful for diabetes research, and how are they applied?
14. What role does insulin play in the regulation of blood glucose levels?
15. How does aging affect the risk of developing type 2 diabetes?
16. Can lifestyle changes reverse type 2 diabetes?

## Appendix - GPT4o Questions

## C. Domain Expert

### C.1. GeneNetwork.org General

Questions with red font did not receive proper answers from GNQA, and were not scored.

1. How do recent advancements in network-based integrative genomics alter our understanding of complex trait architectures?
2. What are the latest methodological improvements in evaluating gene-environment interactions using GeneNetwork.org?
3. How do multi-omics data integration techniques enhance the prediction accuracy of phenotypic traits in GeneNetwork datasets?
4. What are the computational challenges and solutions in analyzing large-scale transcriptomic data within GeneNetwork.org?
5. How has the inclusion of data from diverse populations impacted the generalizability of findings on GeneNetwork.org?
6. What novel insights have been obtained from GeneNetwork.org regarding the genetic basis of psychiatric disorders?
7. How do advancements in machine learning algorithms contribute to the deconvolution of gene expression data in complex tissues?
8. What role do enhancer-promoter interactions play in the regulation of gene networks uncovered through GeneNetwork.org?
9. How can the integration of ATAC-seq data with RNA-seq data on GeneNetwork.org inform about chromatin accessibility and gene regulation?
10. What are the latest strategies for inferring causal relationships within gene networks using data from GeneNetwork.org?
11. How do advancements in single-nucleus RNA sequencing provide more granular insights into cell-type-specific gene expression networks?
12. What impact have recent discoveries in non-coding RNA regulation had on refining gene interaction maps on GeneNetwork.org?
13. How are spatial transcriptomics approaches being integrated into GeneNetwork.org to enhance understanding of tissue architecture and function?
14. How do recent developments in quantitative trait locus (QTL) mapping refine our understanding of gene regulatory variants?”
15. What are the implications of incorporating epigenomic data, such as histone modification maps, into the gene networks on GeneNetwork.org?
16. How do recent findings on 3D genome organization contribute to our understanding of functional genomic networks?
17. What are the potential applications of artificial intelligence in improving the annotation and interpretation of gene networks?
18. How has the study of genetic pleiotropy been advanced by data available on GeneNetwork.org?
19. What novel genetic pathways have been identified in GeneNetwork.org studies related to aging and lifespan?
20. How do polygenic risk scores (PRS) developed using GeneNetwork.org data enhance the prediction and prevention of complex diseases?

### C.2. Aging

1. How do recent single-cell transcriptomics studies enhance our understanding of cellular heterogeneity in aging tissues?”,
2. What are the latest findings on the role of senescence-associated secretory phenotype (SASP) factors in age-related tissue dysfunction?
3. How do age-related changes in chromatin architecture contribute to the decline in cellular function?
4. What insights have been gained from studying the epigenetic reprogramming of aged cells to a more youthful state?
5. How do alterations in the mitochondrial genome and bioenergetics influence the aging process in humans?
6. What are the therapeutic potentials and challenges of targeting the insulin/IGF-1 signaling pathway for extending healthspan and lifespan?
7. How can the integration of proteomics and metabolomics data shed light on age-associated metabolic shifts?
8. What role do long non-coding RNAs (lncRNAs) play in the regulation of aging and age-related diseases?
9. How do recent advancements in CRISPR/Cas9 technology open new avenues for studying and potentially reversing aging?
10. What is the significance of the DNA damage response (DDR) in the context of both replicative and chronological aging?
11. How do age-dependent changes in the immune system, such as immunosenescence, contribute to increased susceptibility to diseases?”,
12. How do advancements in machine learning and artificial intelligence aid in the identification of biomarkers for biological aging?
13. What recent discoveries have been made regarding the impact of systemic factors, such as circulating microvesicles, on aging phenotypes?
14. How do changes in the gut microbiome composition correlate with aging and longevity?
15. What are the key molecular mechanisms through which caloric restriction exerts its lifespan-extending effects across different species?
16. How do oxidative stress and the subsequent accumulation of damaged macromolecules contribute to cellular aging?
17. How are extracellular matrix remodeling and tissue stiffness implicated in the aging process?
18. How do recent developments in autophagy research contribute to our understanding of its role in aging and longevity?
19. What are the implications of age-related shifts in stem cell niche composition and function for tissue regeneration capacity?
20. How do cross-links and advanced glycation end-products (AGEs) accumulation affect the structural integrity and function of aging tissues?

### C.3. Diabetes

1. How do recent advancements in multi-omics approaches, including proteomics and metabolomics, contribute to our understanding of Type 2 diabetes pathogenesis?
2. What novel diabetic loci have been identified through the latest meta-analyses of large-scale genome-wide association studies (GWAS)?
3. How do epigenetic modifications, such as DNA methylation and histone modification, influence the expression of diabetes-related genes?
4. Can you elaborate on the role of the gut microbiome in modulating host genetic predispositions to diabetes?
5. How effective are machine learning algorithms in integrating genomic data to predict individual risk and progression of diabetes?
6. What are the implications of recent findings on the role of long non-coding RNAs (lncRNAs) in the regulation of insulin secretion and sensitivity?
7. How do post-translational modifications of proteins affect key signaling pathways involved in glucose homeostasis?
8. What insights have been gained from studying the genetic basis of syndromic forms of diabetes, such as Wolfram Syndrome and Alström Syndrome?
9. How do genetic and epigenetic differences between monozygotic twins discordant for diabetes inform our understanding of its etiology?
10. What potential therapeutic targets have been identified through recent studies on the interaction between genetic variants and environmental factors in diabetes development?
11. How do rare variants identified through whole-genome sequencing contribute to the heritability of Type 2 diabetes?
12. What are the latest findings on the role of non-coding RNAs in the pathogenesis of diabetes?
13. How does the interaction between multiple polygenic risk scores (PRS) improve the prediction of Type 1 and Type 2 diabetes?
14. What are the mechanistic insights into the beta-cell failure pathways gleaned from recent single-cell RNA-sequencing studies?
15. How does the epigenetic landscape of key metabolic tissues change in diabetic versus non-diabetic individuals?
16. What recent advancements have been made in leveraging CRISPR-based approaches to correct monogenic forms of diabetes in vivo?
17. How do genome-wide association studies (GWAS) integrate with multi-omics data to elucidate the complex genetic architectures of diabetes?
18. What is the impact of genomic imprinting on the susceptibility and progression of diabetes?
19. How do longitudinal genomics studies help in understanding gene-environment interactions in diabetes onset and management?
20. How have recent integrative genomics approaches, such as the use of single-cell RNA sequencing combined with epigenomic profiling, advanced our understanding of cellular heterogeneity and gene regulatory networks in pancreatic beta cells under diabetic conditions?

## D. Citizen Scientist

### D.1. GeneNetwork.org General

Questions with red font did not receive proper answers from GNQA, and were not scored.

1. What is GeneNetwork.org, and how does it help scientists understand genetics?
2. How do researchers use GeneNetwork.org to study diseases?
3. What can GeneNetwork.org tell us about how genes interact with each other?
4. How does GeneNetwork.org help in finding the genetic causes of common diseases?
5. Can GeneNetwork.org predict my risk of developing certain health conditions based on my genes?
6. How does GeneNetwork.org make use of data from different populations around the world?
7. What kinds of genetic data are available on GeneNetwork.org?”
8. How do scientists use GeneNetwork.org to study differences in gene expression?
9. Can GeneNetwork.org be used to learn about genetic influences on behavior?
10. What role does GeneNetwork.org play in personalized medicine?
11. How does the information on GeneNetwork.org help in developing new treatments for diseases?
12. What is a gene network, and why is it important for understanding genetics?
13. How do researchers identify which genes are important for certain traits using GeneNetwork.org?
14. How can GeneNetwork.org help in understanding complex traits like height or intelligence?
15. Are there any known genetic mutations that cause premature aging?
16. What are the practical applications of the research done through GeneNetwork.org?
17. How can I access and use the data available on GeneNetwork.org?
18. What are some recent discoveries made using GeneNetwork.org?
19. How do scientists ensure the accuracy of the data on GeneNetwork.org?
20. What’s the difference between looking at one gene and studying a whole gene network?

### D.2. Aging

1. 2. What are the main genetic factors that influence aging?
2. How do genes affect the aging process in humans?
3. What lifestyle choices can help slow down genetic aging?
4. How do scientists study the genetics of aging in animals?
5. Are there specific genes that have been linked to longer lifespans?
6. How do telomeres affect the aging process?
7. What role does DNA repair play in aging?
8. Can genetic research lead to treatments that slow down aging?
9. How does mitochondrial DNA influence aging?
10. Are there any known genetic mutations that cause premature aging?
11. What recent discoveries have been made about the genetics of aging?
12. How do epigenetic changes affect aging?
13. What is the role of the gene FOXO3 in longevity?
14. How does the environment interact with genes to influence aging?
15. What are senescent cells and how do they contribute to aging?
16. Are there any known lifestyle interventions that can positively impact genes related to aging?
17. What is the ‘epigenetic clock,’ and how is it used in aging research?
18. How do researchers use model organisms like yeast or worms to study human aging?
19. Are there any promising anti-aging therapies being developed based on genetic research?
20. How do caloric restriction and diet impact the genetics of aging?

### D.3. Diabetes

1. How do genetic mutations in the insulin gene affect glucose metabolism?
2. What are the most common genetic loci associated with an increased risk of Type 2 diabetes?
3. How does genome-wide association studies (GWAS) help in identifying diabetes-related genes?
4. What is the role of the HLA region in the genetic predisposition to Type 1 diabetes?
5. How do genetic differences contribute to variations in diabetes prevalence among different populations?
6. What is the function of the PPAR-gamma gene in diabetes, and how do its variants impact the disease?
7. How can CRISPR/Cas9 technology be used to study or treat genetic forms of diabetes?
8. What is the significance of genetic polymorphisms in the GLUT4 gene for Type 2 diabetes?
9. How do microRNAs regulate gene expression related to diabetes?
10. What insights have been gained from studying the genetic basis of MODY (Maturity Onset Diabetes of the Young)?
11. What genes are most commonly associated with an increased risk of developing diabetes?
12. How can genetic testing help predict a person’s risk for diabetes?
13. What role do family genetics play in the likelihood of getting diabetes?
14. Can lifestyle changes affect genetic risk factors for diabetes?
15. What recent breakthroughs have been made in understanding the genetic causes of diabetes?
16. How do genes influence how our bodies respond to sugar and insulin?
17. Are there specific genetic markers that can indicate a higher risk for Type 1 versus Type 2 diabetes?
18. How can new gene therapies potentially cure or treat diabetes?
19. What is the difference between monogenic and polygenic diabetes?
20. How does studying the DNA of people with diabetes help scientists find better treatments or cures?

## RAGAS Evaluation Equations

The equations are taken from RAGAS documentation.

### Faithfulness

This measures the factual consistency of the generated answer against the given context. It is calculated from answer and retrieved context. The answer is scaled to (0,1) range. Higher the better.

The generated answer is regarded as faithful if all the claims made in the answer can be inferred from the given context. To calculate this, a set of claims from the generated answer is first identified. Then each of these claims is cross-checked with the given context to determine if it can be inferred from the context. The faithfulness score is given by:

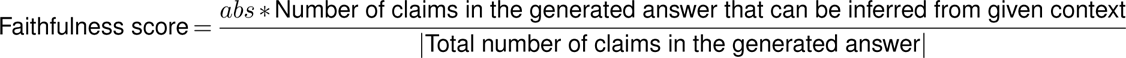

### Context Utilization

Context utilization is like a reference free version of context_precision metrics. Context utilization is a metric that evaluates whether all of the answer relevant items present in the contexts are ranked higher or not. Ideally all the relevant chunks must appear at the top ranks. This metric is computed using the question, answer and the contexts, with values ranging between 0 and 1, where higher scores indicate better precision.

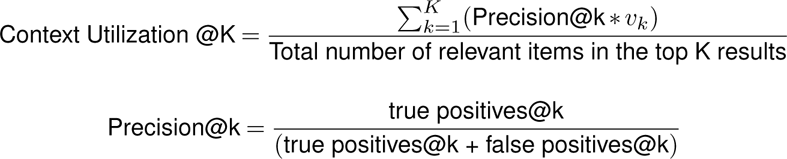

Where *K* is the total number of chunks in contexts and *v_k_* ∈ {0, 1} is the relevance indicator at rank *k*.

### Answer Relevance

The evaluation metric, Answer Relevancy, focuses on assessing how pertinent the generated answer is to the given prompt. A lower score is assigned to answers that are incomplete or contain redundant information and higher scores indicate better relevancy. This metric is computed using the question, the context and the answer.

The Answer Relevancy is defined as the mean cosine similarity of the original question to a number of artifical questions, which where generated (reverse engineered) based on the answer:

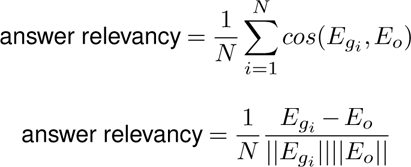

Where:

- *E_g_i__* is the embedding of the generated question *i*.
- *E_o_* is the embedding of the original question.
- *N* is the number of generated questions, the default of which is 3.

Please note, that even though in practice the score will range between 0 and 1 most of the time, this is not mathematically guaranteed, due to the nature of the cosine similarity ranging from −1 to 1.

### Context Relevance

Content for this sub section comes from **Shahul:2023**, the author of RAGAS: Automated Evaluation of Retrieval augmented generation [**Shahul:2023**].

The context *c*(*q*) is considered relevant to the extent that it exclusively contains information that is needed to answer the question. In particular, this metric aims to penalise the inclusion of redundant information. To estimate context relevance, given a question *q* and its context *c*(*q*), the LLM extracts a subset of sentences, *S_ext_*, from *c*(*q*) that are crucial to answer *q*, using the following prompt:

> *Please extract relevant sentences from the provided context that can potentially help answer the following question. If no relevant sentences are found, or if you believe the question cannot be answered from the given context, return the phrase “Insufficient Information”. While extracting candidate sentences you’re not allowed to make any changes to sentences from given context*.

The context relevance score is then computed as:

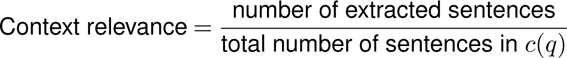

## Notes

https://git.genenetwork.org/gn-ai/tree/README.md

### Summary of Updates

In the original write up the author list is different than how the list was input in the system. It was assumed the author list from the pdf would translate to the site and it did not; hence, the author list was reordered on the 1st page beneath the title.

